# Divergent genetic mechanism leads to spiny hair in rodents

**DOI:** 10.1101/267385

**Authors:** Gislene L. Gonçalves, Renan Maestri, Gilson R. P. Moreira, Marly A. M. Jacobi, Thales R. O. Freitas, Hopi E. Hoekstra

## Abstract

Spines, or modified hairs, have evolved multiple times in mammals, particularly in rodents. In this study, we investigated the evolution of spines in six rodent families. We first measured and compared the morphology and physical properties of hairs between paired spiny and non-spiny sister lineages. We found two distinct hair morphologies had repeatedly evolved in spiny rodents: hairs with a grooved cross-section and a second near cylindrical form. Compared to the ancestral elliptical-shaped hairs, spiny hairs had higher tension and stiffness, and overall, hairs with similar morphology had similar functional properties. To examine the genetic basis of this convergent evolution, we tested whether a single amino acid change (V370A) in the *Ecdysoplasin A receptor (Edar)* gene is associated with spiny hair, as this substitution causes thicker and straighter hair in East Asian human populations. We found that most mammals have the common amino acid valine at position 370, but two species, the kangaroo rat (non-spiny) and spiny pocket mouse (spiny), have an isoleucine. Importantly, none of the variants we identified are associated with differences in rodent hair morphology. Thus, the specific *Edar* mutation associated with variation in human hair does not seem to play a role in modifying hairs in wild rodents, suggesting that different mutations in *Edar* and/or other genes are responsible for variation in the spiny hair phenotypes we observed within rodents.

## Introduction

Hair is a defining trait of mammals yet can be highly differentiated among species, primarily with respect to density and morphology. For example, sea otters (*Enhydra lutris*) have the densest fur of any mammal, with up to 140,000 hairs/cm^2^, more than on an entire typical human head [1]. A second extreme example is the modification of hairs into spines, which have evolved multiple times in mammals, including echidnas (Tachiglossidae), tenrecs (Tenrecidae), hedgehogs (Erinaceidae) and rodents (Hystricidae and Erethizontidae) [2]. In addition, a second type of spine, defined as aristiform (i.e., bristle shaped) hair [3], has evolved independently at least five times within Rodentia [2, 4, 5].

Mammalian coats are comprised of four different types of hair each with a distinct morphology: guard, awl, auchene and zigzag hairs [6]. These four types can be distinguished mainly based on length, number of medulla cells and the presence of bends in the hair shaft [7]. In general, spines and aristiform hairs are considered to be a modified hair, sharing similarities to guard hairs [2, 14]. However, aristiform hairs of the spiny mouse (*Acomys*) likely represent enlarged awl hairs, based on a developmental comparison with a mouse model [8].

Yet, despite the many descriptions of the structural features of spines, and their use in taxonomic studies [2, 4, 9–15], the evolutionary patterns and processes of morphological variation in these modified hairs are poorly understood. Similarly, the identification of the genes and mutations underlying variation in hair structure is still nascent.

The role of the *Ectodysplasin A receptor* (*Edar*) gene, homolog to *Downless* in mouse [16], has been implicated in the morphological differentiation of hair and other
ectodermic appendages in humans [17–20]. This gene encodes a novel member of the tumor necrosis factor (TNF) receptor family that include an extracellular cysteine-rich fold, a single transmembrane region, and a death homology domain close to the C terminus [18, 21]. This transmembrane protein is a receptor for the soluble ligand *ectodysplasin A* (*Eda*) and can activate the nuclear factor-kappa B, JNK, and caspase-independent cell death pathways [22]. Mutations that inhibit this developmental pathway prevent the formation of hair follicles in mice and humans [23–25], indicating the involvement of this gene in the development of hair, teeth, and other ectodermal derivatives [26].

In humans, a nonsynonymous single nucleotide polymorphism (SNP) in the *EDAR* gene, T^1540^C (rs3827760), causes a Val^370^Ala amino acid change, which is associated with variation in hair structure in East Asian populations [27, 28]. Experiments in cultured cells have shown that this specific mutation enhances *Edar* signaling *in vitro*, which in turn alters multiple aspects of mouse hair morphology including straighter fibers, increased diameter, and more cylindrical form compared to hair of either European or African origin [29]. A knock-in mouse model reinforced that, as in humans, hair thickness is increased in mice carrying 370A [19]. Moreover, evidence from clinical genetics shows that EDAR 370A causes increased function, as it suppresses the effects of a hypomorphic EDA mutation [30].

Interestingly, the *Edar* mutant mouse [26] has pelage that bears remarkable resemblance to that of wild mice with aristiform hairs like, for example, the Southern African spiny mouse (*Acomys spinosissimus*). This gross-level phenotypic similarity led us to ask whether similar evolutionary pathways – at both the phenotypic and genetic levels – are responsible for the East Asian human hair phenotype and the spiny hairs observed in many rodent clades. *Edar* is a strong candidate for explaining variation in hair morphology, thus we have focused on this gene and the Val^370^Ala mutation specifically.

In this study, we investigate the hair phenotype of all six families of rodents that have spiny hairs (Table 1). First, we characterize their specific phenotype by comparing types of hairs (“spiny” and “control” guard hair) between closely related species, focusing on variation in both morphology such as size and shape as well as physical properties. Second, we test for an association between genotype at the homologous amino acid position 370 in *Edar* and hair morphology. Together, these data allow us to determine if spiny hairs have evolved through similar or different evolutionary pathways in humans and rodents.

**Table 1.**
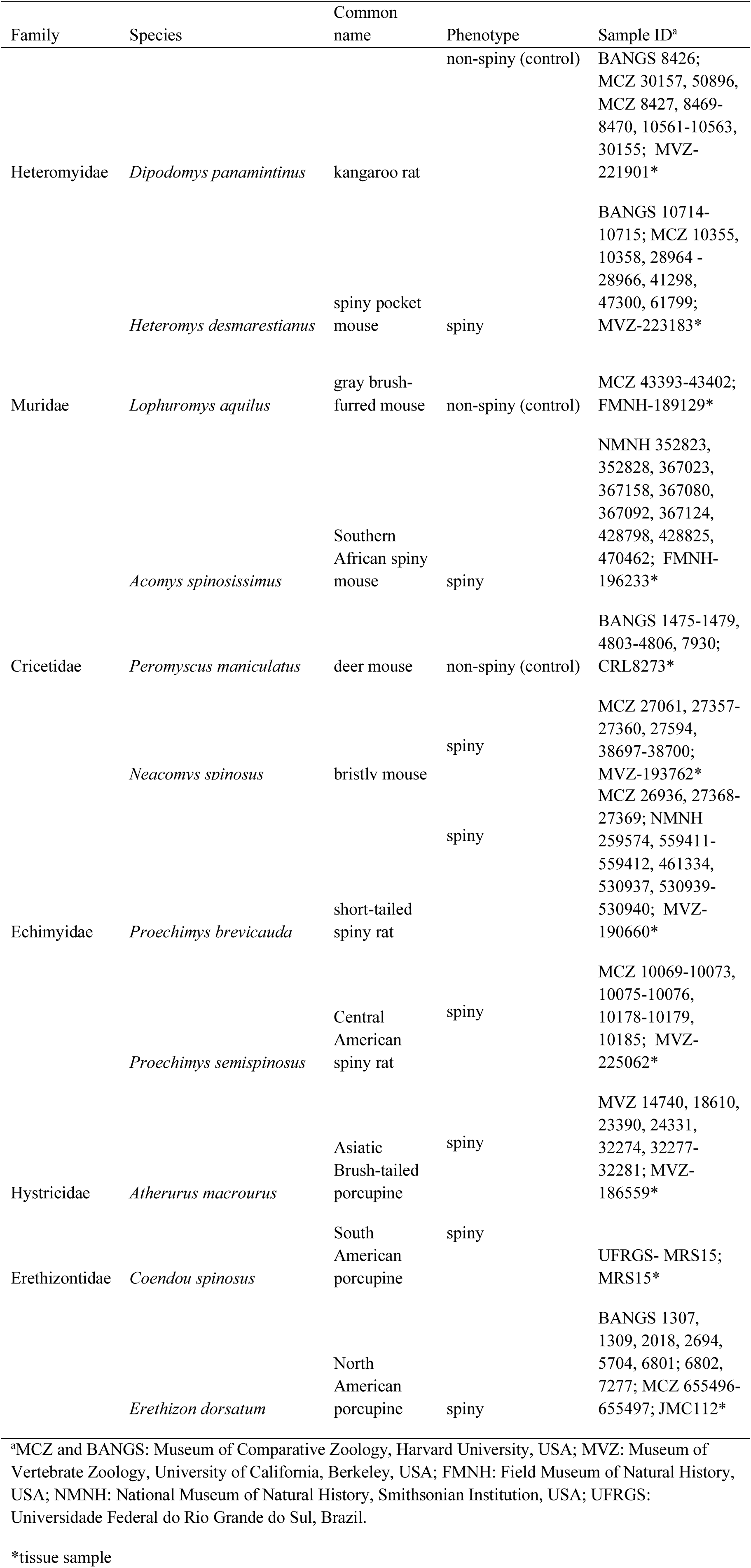
Rodents specimens analyzed in this study.

## Material and Methods

We analyzed eight species that represent all six extant families of rodents that include at least one species with spiny hair: Spiny pocket mouse (*Heteromys desmarestianus*) [Heteromyidae], Southern African spiny mouse (*Acomys spinosissimus*) [Muridae], Bristly mouse (*Neacomys spinosus*) [Cricetidae], Short-tailed spiny rat (*Proechimys brevicauda*) and Central American spiny rat (*Proechimys semispinosus*) [Echimyidae], Asiatic brush-tailed porcupine (*Atherurus macrourus*) [Hystricidae], South American porcupine (*Sphiggurus villosus*) and North American porcupine (*Erethizon dorsatum*) [Erethizontidae] (Table 1). Within these six rodent families, we next identified a closely related non-spiny taxa: Kangaroo rat (*Dipodomys panamintinus*) [Heteromyidae], Gray brush-furred mouse (*Lophuromys aquilus*) [Muridae] and Deer mouse (*Peromyscus maniculatus*) [Cricetidae] to control for the possible confounding effects of shared evolutionary history. Tissue samples from these 11 species were provided by the following museum collections: Museum of Comparative Zoology [MCZ], Harvard University; Museum of Vertebrate Zoology [MVZ], University of California, Berkeley; Field Museum of Natural History [FMNH]; National Museum of Natural History [NMNH], Smithsonian Institution; and Federal University of Rio Grande do Sul [UFRGS].

### Morphometric analysis

For morphological analyses, we focused on modified hairs (i.e. spines and aristiforms) in the focal species and non-modified guard hairs in the phylogenetically-paired control species. We defined guard hairs as those that were the longest, had two medulla cells and did not have any bends in the hair shaft.

Of the 11 taxa surveyed, we used a total of 10 specimens per species for the morphological and molecular analyses. All hairs were plucked from the lower left hip area to minimize differences due to pelage variation across the body. Hairs were gently removed with a pair of fine-tipped, self-closing forceps that caused no visible damage to the sample.

We first photographed each hair on a flat surface, and then cross-sectioned the hair with a scalpel and mounted the resultant sections on glass slides. We captured digital images, using a Sony Cyber-shot DSC-H10 digital camera attached to a stereomicroscope Leica M125 at the thickest point of each hair. We measured linear parameters, such as hair length, width, ellipticity and concavity (Fig 1). Cross-sectional ellipticity (*sensu* [29]) was calculated as a ratio of between width and thickness (= d/c), and concavity as the ratio between the depth of the dorsal groove (= zero, when absent) and cross section width (= b/d). We performed all hair measurements on digital images using AxioVision microscopy software (Zeiss).

**Fig 1.**
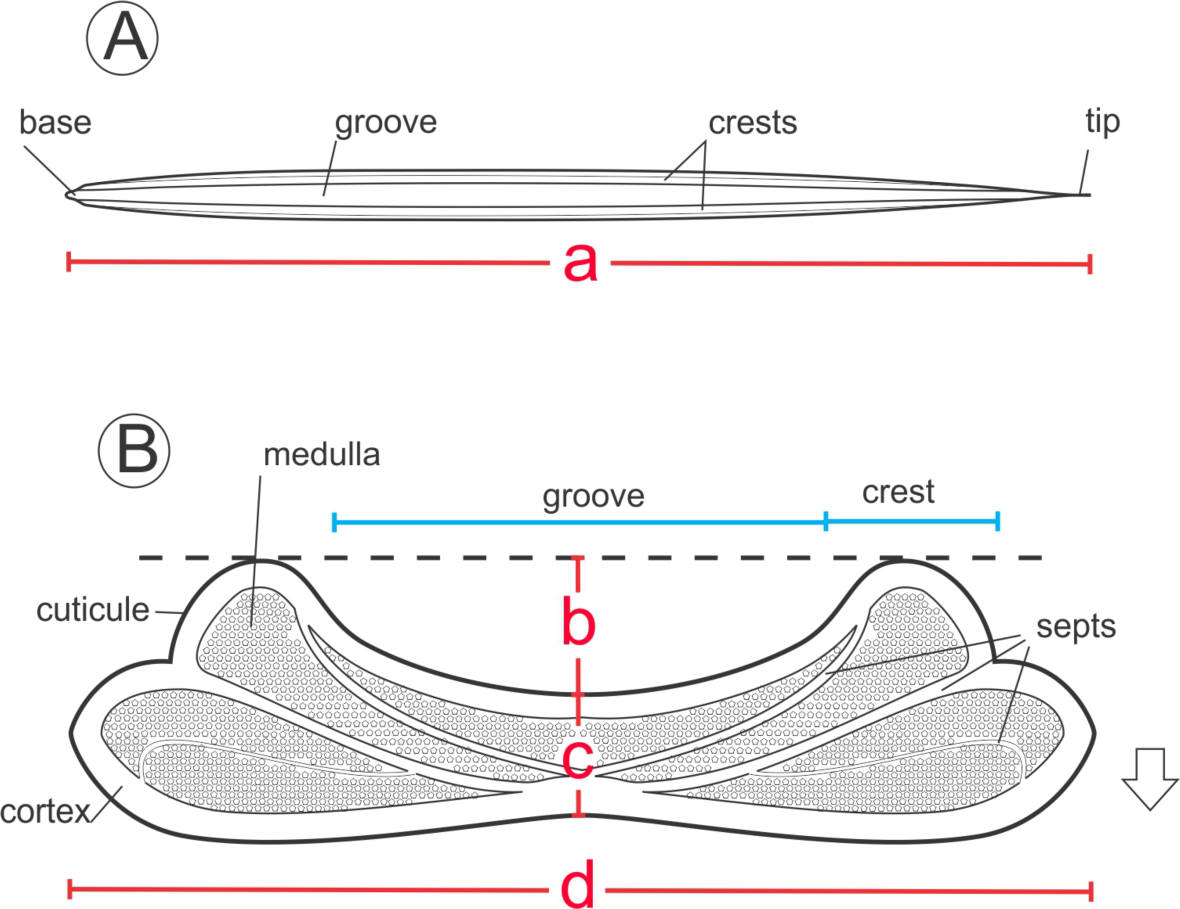
Schematic representation of a spiny hair showing (A) dorsal view and (B) cross section, with structural components. Measurements include: a = length, b = depth of groove, c = thickness, d = width. Arrow indicates the dorsal-ventral direction.

To measure the cross-sectional dimension of hairs using scanning electron microscopy, additional hairs were laid on a hard surface and cut transversally into 1-cm sections with a razor blade. Then, we mounted these sections with double-sided tape on metal stubs, coated with gold in a Bal-tec^®^ SCD050 sputter coater to examine and photograph them using a JEOL JSM6060 scanning electron microscope at the Centro de Microscopia Eletrônica (CME) of UFRGS. To describe the ultrastructure of hair surface, we employed nomenclature previously used [2], and for the shape of cross sections, we followed Teerink [3].

To detect evolutionary convergence and divergence, we visualized differences in hair shape among species using linear measurements to conduct a principal component analysis (PCA). A phylogenetic tree was projected onto the form space of the two first principal components, creating a phylomorphospace [31, 32]. Internal forms at nodes were reconstructed by squared-change parsimony. The phylomorphospace was created using phytools [33] in R [34].

### Hair tension analysis

To generate hair tension values (σ) as function of the relative deformation (ε), we used a universal testing machine EMIC DL 5000/10000 with a strain rate of 1mm/min. First, to calculate the initial transverse area, the depth, and thickness of the flat hair samples and the radius for the cylindrical samples, we used a manual digital pachymeter with an accuracy of 0.01 mm. We took a sample from the middle section each individual hair. Next, we glued the distal end of each hair with water putty (Durepox) to a small pieces of sand-paper (20 × 20 mm). This process increased friction, thus allowing stable anchoring of samples to the tension grips of the machine. We calculated the tension, σ=N/A_0_, expressed by MPa (N/mm^2^), from the force necessary to deform the sample until its rupture expressed in Newtons (N) divided by the transversal initial area of the sample, A_0_ (A_0_ = depth x thickness for the concave hair and A_0_ = π.r^2^ for the cylindrical sample). Specifically, we used the sample’s initial length (L_0_) and load cell of 10mm and 500N for the thicker hairs (e.g., porcupines), and 5 mm and 50N for the finer hairs. We also measured the absolute relative deformation, e, that is equal to (L-L_0_)/L_0_, in which L = the length of the sample at testing, and L_0_ = the initial length of the sample. The tensile strength and the deformation are sensitive to macroscopic defects at the surface of the sample. Consequently, from the initial ten samples measured for each species, we selected the best (i.e., least worn with no visible defects) specimens to use in an interspecific comparison, which resulted in only small variation in the physical properties of hairs within species. Finally, by plotting tension as function of deformation, we generated the tensile curves for each species.

In addition, to estimate hair stiffness, we measured the Young’s modulus, E = σ/ε in the linear (or elastic) deformation region. This is a measure of tensile elasticity, or the tendency of an object to deform along an axis when opposing forces are applied along that axis. We calculated Young’s modulus as the slope of the curve in its initial linear region (up to 3% deformation for all species). We then generated a bar plot comparing values of Young’s modulus in a phylogenetic context using the phytools package in R.

### Screening for EDAR V370A polymorphism

For our genetic analyses, we extracted total genomic DNA from tissue samples (muscle) preserved in DMSO using Blood and Tissue Kit (Qiagen) following manufacturer instructions. DNA was then stored at -20°C. Degenerate primers were designed to amplify the final exon of *Edar* (Table 2), which encompass the variant V370A in humans. Because this is a highly conserved region [16], and several mutations were previously demonstrated to be involved in human clinical pathologies [35], substitutions in this region may be functionally relevant.

**Table 2.**
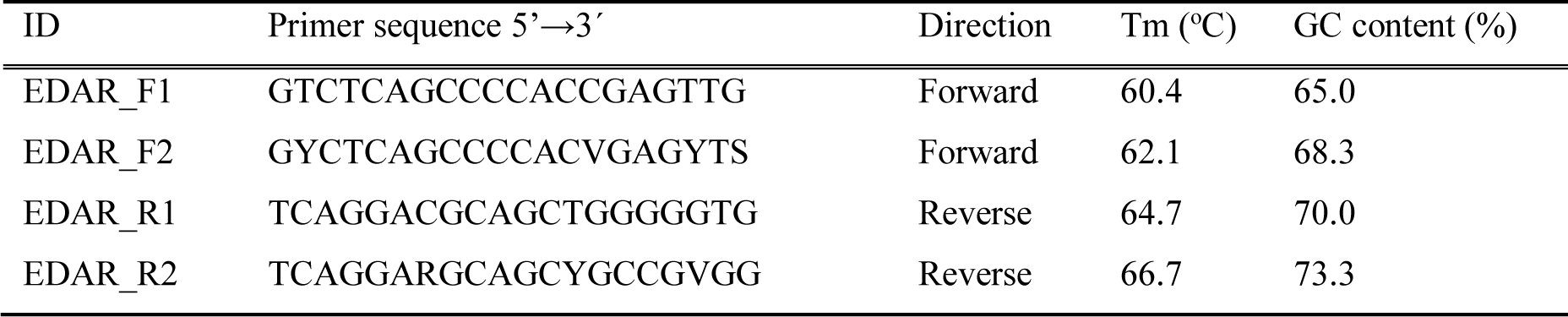
Primers designed to amplify a 275-bp region of *Edar* gene that encompasses the death domain and the V370A variant.

For the amplification reaction, we performed a touchdown PCR (i.e., decreasing annealing temperatures from 60 °C to 50 °C) with AmpliTaq Gold (Perkin Elmer) and 1.8 mM MgCl_2_. We used primer pair F1+ R2 to amplify species in the Cricetidae, Heteromyidae and Muridae lineages and F2+R1 for Echimyidae, Erethizontidae and Hystricidae. We next verified the resultant PCR products in agarose gel stained with SybrSafe (Invitrogen), extracted the bands, and purified them using QIAquick Gel Extraction kit (Invitrogen). Using the same primer pairs, we next sequenced the fragments in both directions using BigDye chemistry in an automated sequencer ABI3730XL (Applied Biosystems). We next aligned all the sequence data using Codon Code Aligner (CodonCode Corp.).

To infer phylogenetic relationships among the rodents surveyed, we used a mitochondrial DNA marker (*Cytochrome b* [*Cytb*]). For seven species, *Cytb* sequences were already available in GenBank (Supplementary Material); for three others, we sequenced a fragment of the gene (ca. 800 bp) using primers MVZ05 and MVZ16 with conditions described in Smith & Patton [36]. We constructed a phylogenetic tree using the maximum likelihood method with heuristic search option, tree bisection–reconnection, and an initial neighbor joining clustering. Branch support was estimated by 1000 bootstrap replicates using a heuristic search of nearest-neighbor interchange.

## Results

### Variation in hair morphology

We uncovered substantial qualitative and quantitative variation in hair morphology among rodent species. While we found a gradient of shape in the cross sections of hairs, we could group the morphologies into three major forms: elliptical, grooved, and circular (Fig 2), the latter two of which we considered “spiny” phenotypes. The non-spiny guard hairs of the three control species (i.e., kangaroo rat, gray brush-furred rat and deer mouse) were all elliptical and did not have a differentiated lateral groove or ridges (Fig 2A-C). The grooved form varied dramatically, including spines with different degree of hardness. The spiny hairs present in the bristly mouse, spiny pocket mouse and Southern African spiny mouse are characterized by a pattern of crests on the dorsal surface and a decrease in the ventral curvature (Fig 2D-I). Unlike the grooved hairs, medulla of the circular form in new world porcupines (Erethizontidae) have several uniform alveolar cells of small size that fill the entire lumen without septa and are associated with an interleaved cortex of reduced thickness (Fig 2J, K). Thus, spiny hairs, compared to control hairs, all share a dorsal groove associated with longitudinal side ridges, with one exception, the New World porcupines (Erethizontidae) (Fig 2D-I).

**Fig 2.**
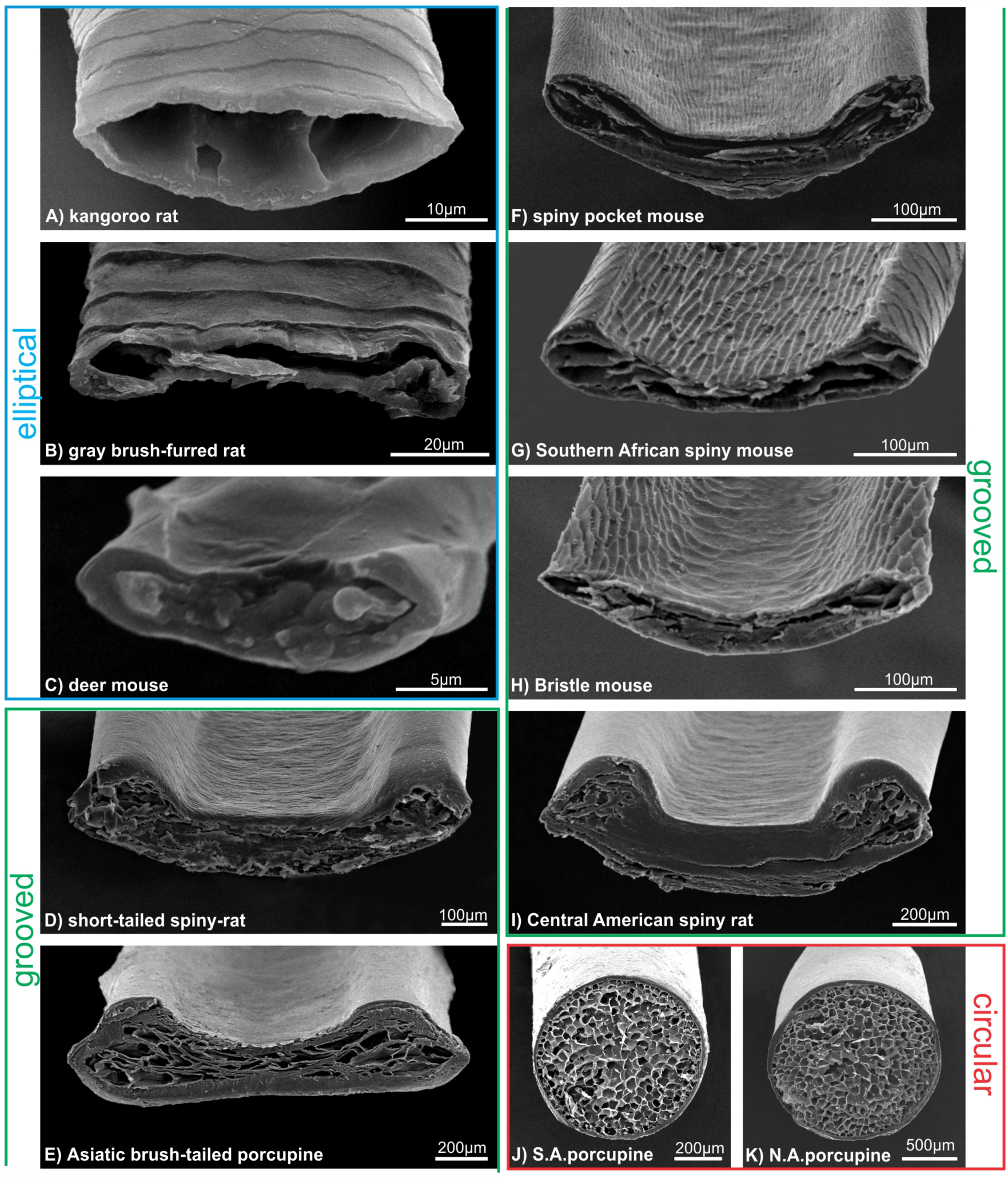
Scanning electron micrographs of cross sections of guard hairs, indicating the three major morphologies observed: elliptical (A-C), grooved (D-H) and circular (D-K).

The grooved hair type was particularly common in our data set (6 of 11 taxa) and showed markedly differences among lineages. These hairs differed in length, width and thickness (Fig 3). First, the Asiatic brush-tailed porcupine is the most distinct, having the widest hairs with a large area. Second, the short-tailed and Central American spiny rats had similar hair phenotype to each other and had intermediate values – between the brush tailed porcupine and all other species – of length, width, thickness and area. Third, the spiny pocket mouse, Southern African spiny mouse and bristle mouse showed low values in length, width and thickness. Therefore, a gradient from harder (Asiatic brush-tailed porcupine) to softer (the spiny pocket mouse, Southern African spiny mouse and bristle mouse) hairs was evident within the grooved hair type. We also found a larger perimeter and cross-sectional area in softer spines compared to harder spines.

**Fig 3.**
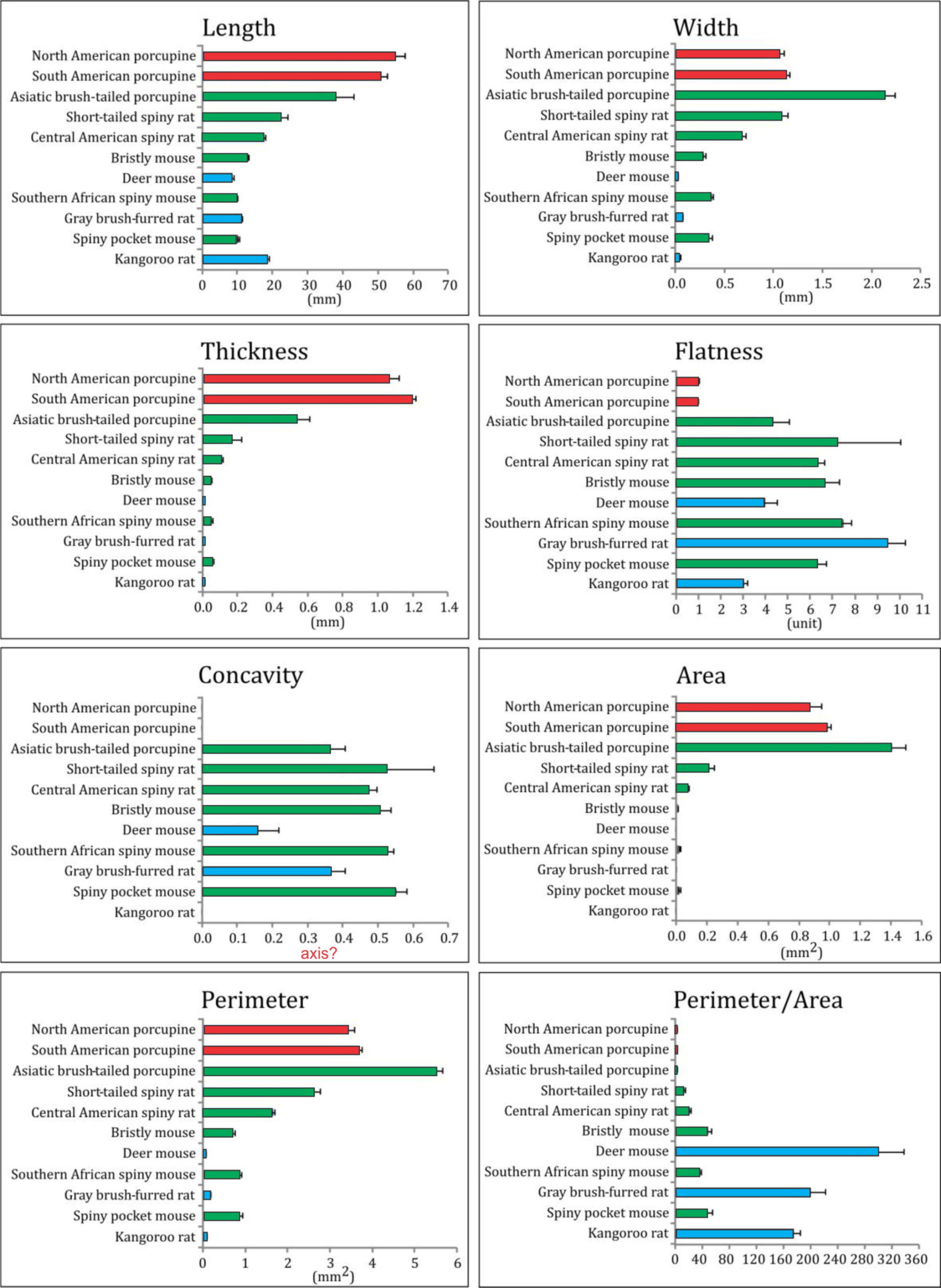
Variation in guard hairs and spiny hairs among rodent species. Color indicates the cross-sectional morphology (as in Fig 1). Sample size=10 specimens/species. Mean ± SE is provided.

We did not, however, find a clear pattern in the degree of flatness across phenotypes (Fig 3). The extent of concavity was similar among hairs with different degrees of hardness, but was much lower in the non-modified guard hairs of the control species (deer mouse, gray brush-tailed rat and kangaroo rat).

The unique, hard spines observed in the two species of New World porcupines (Erethizontidae) were both longer and thicker than any other taxa sampled (Fig 3). The cylindrical shape of these spines (observed in cross section; Fig 2J, K) resulted in a small perimeter/area ratio. Moreover, this cylindrical shape, and lack of groove and ridges, resulted in a flatness score near one and a concavity value of zero (Fig 3).

To summarize the variation in hair morphology in a phylogenetic context, we plotted each species’ mean hair shape in phylomorphospace. We found a remarkable degree of convergence in hair form among rodents with elliptical and grooved hair phenotypes (Fig 4). Overall, principle component 1 (PC1), which captures 72% of the variation, largely separates elliptical and circular hair shapes, and PC2 (22% of the variation) separates the grooved hairs. Pairs of related species (within the families Heteromyidae, Cricetidae and Muridae) are separated in morphospace, reflecting their divergence in hair type despite their phylogenetic similarity. In other words, hair morphology does not closely reflect phylogenetic relationships, and other ecological factors, for example, are likely driving the observed morphological convergence.

**Fig 4.**
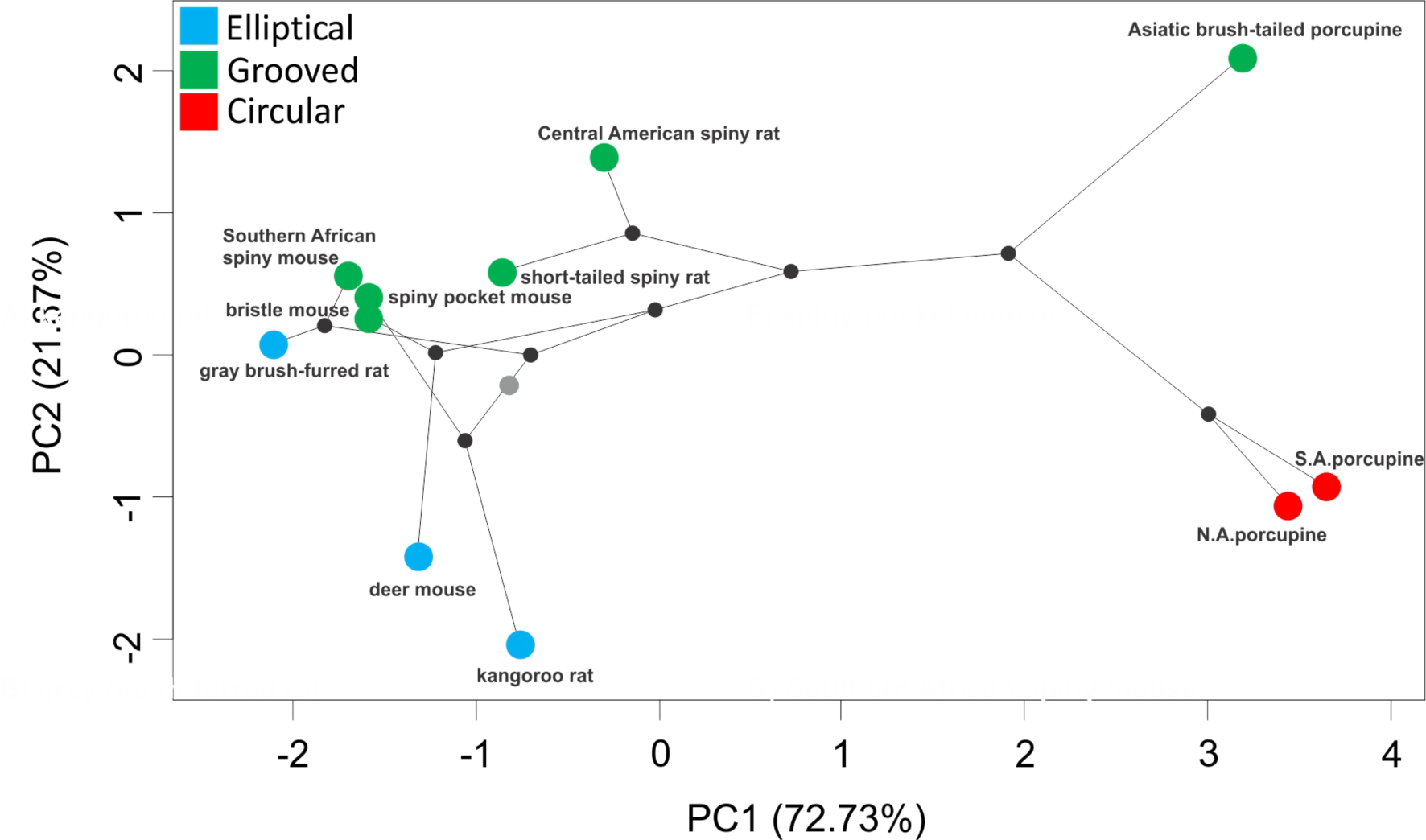
A phylomorphospace plot of hair morphology for 11 rodent species. Color indicates cross-sectional morphology: blue, elliptical; green, grooved; red, circular. A phylogeny based on cyt-*b* sequences is depicted on the morphological space. Nodes are represented by black dots, and the root by a gray dot.

### Hair tension and deformation

To quantify how differences in hair morphology affect functional variation, we measured both hair tension and force as hairs were subjected to deformation. First, we found that the tension curves of non-spiny control species with elliptical hairs are distinct from the grooved and circular hairs (Fig 5A). For elliptical hairs, tension can reach high values (e.g. up to 140 MPa) for the first 2-5% of deformation before the tension, σ, stabilized (around 20 MPa) and then, ultimately, the hair breaks. These elliptical hairs also show low mechanical resistance (Fig 5B). Grooved hair all had similar tensile curves, in which tension values increased in a roughly linear fashion for the first ~10% of hair deformation then slowly stabilized (Fig 5A). Their maximum resistance, the peak in the curve after stabilization, was near the limit of deformation for most of these grooved species. Tension values were lower for the Asiatic porcupine and both the short-tailed and Central American spiny rats compared to the relative higher tension values of the grooved hair (Fig 5A). Finally, the American porcupines (Erethizontidae) with circular hair shape have curves in which, after a shallow upward slope, tension remained constant over increasing deformation (Fig 5A). Moreover, in our dataset, these spines also require the highest absolute force needed to deform them (Fig 5B). Thus, hair shape appears to affect both its tensile strength and the force required to deform the hair.

**Fig 5.**
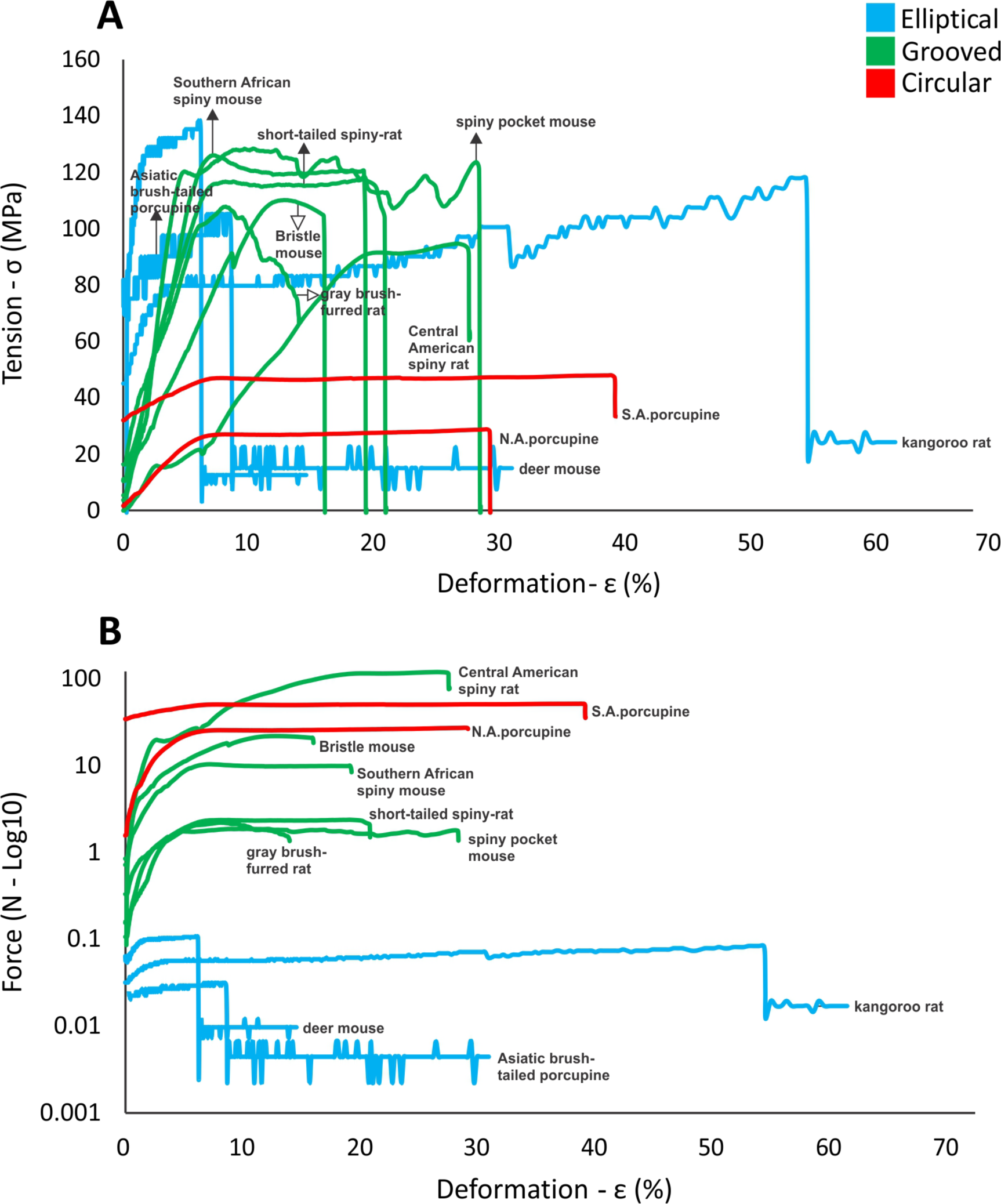
Deformation curves showing (A) tension and (B) force for hairs from 11 rodent species (labels correspond to species in Fig. 2).

To measure the stiffness of hairs, we calculated the Young’s elastic modulus for hairs of each species. We found that elliptical hairs, of the non-spiny control species, had the highest values, indicating they are less stiff, intermediate values for the grooved hairs, and the lowest values for the circular hairs of the porcupines (Fig S1). One notable exception was the Asiatic porcupine, whose hairs had stiffness values similar to the New World porcupines, despite having grooved hair morphology. Overall, these data reinforce a pattern of convergence, in which similar hair morphologies have similar functional measurements.

### EDAR variants

To test for an association between hair phenotype and genotype, we sequenced 275 bp of the *Edar* gene in all 11 rodent species. Importantly, this segment spans exon 11 which contains the Val^370^Ala mutation, previously implicated in hair differentiation in East Asian populations of humans. When comparing this region across these species, we observed a number of nucleotide substitutions (54 variable sites; Table S1); however most did not result in amino acid change. In fact, this region showed 98% amino acid conservation among these taxa (Fig 6). Most notably, the 370A was not present in any rodent lineages, either spiny or non-spiny controls (Fig 6A). The first and third position of codon 370, however, did vary among species (whereas the second remained conserved), which resulted in one missense mutation (Val^370^Iso), present in both the kangaroo rat (non-spiny phenotype) and the spiny pocket mouse, and thus not associated with hair phenotype (Fisher’s Exact test, p=0.49; Fig 6B). A second, conservative change at amino acid position 422 (Leu^422^Val) was observed in only the tree rats, two closely related species. Thus, we found no obvious correlation between hair phenotype and the candidate amino acid change at position 370, and no new changes associated with hair morphology at other positions in the death domain of EDAR.

**Fig 6.**
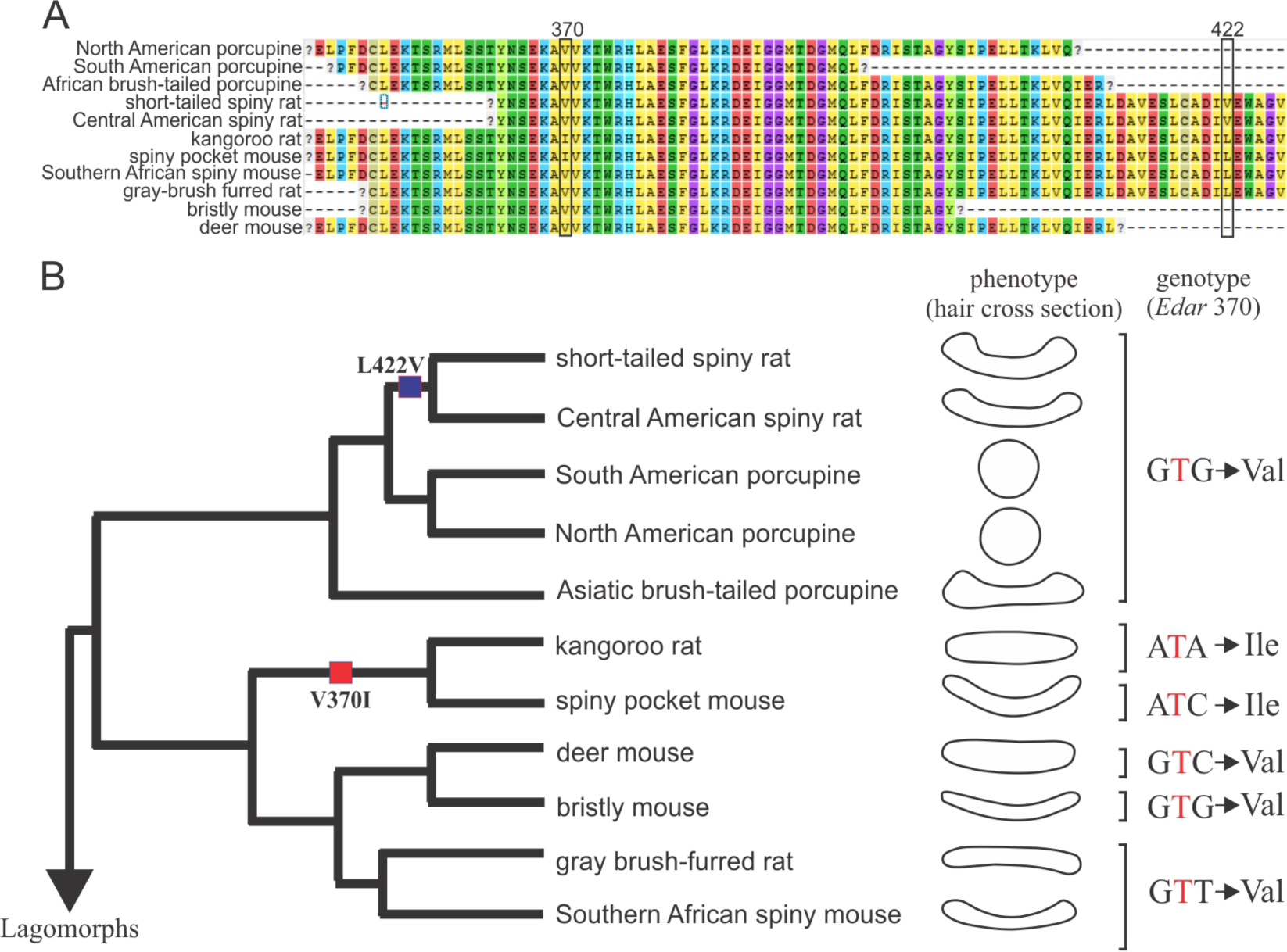
Amino acid sequence alignment of EDAR protein, hair morphology and variability in 11 rodents. A) Variability along 275 bp of the last exon in *Edar* gene, indicating the two missense changes by squares. B) Bayesian phylogenetic tree reconstructed based on partial sequences of the cytochrome b gene. Two amino acid substitutions, V^370^I and L^422^V, are mapped onto phylogeny. Hair morphology of each species is depicted by a cartoon of its cross section. Genotypes at the amino acid position 370 are indicated by the triplet of nucleotides.

## Discussion

In this study, we show that, at the level of morphology, two evolutionary pathways exist to make a hair spiny in wild rodents. Specifically, in the bristle mouse, spiny pocket mouse, Southern African spiny mouse, tree rats and Asiatic brush-tailed porcupine, a convergent pattern emerges: from non-spiny elliptical hairs, the hairs of
these rodent species not only in increase in size (length, width and thickness) but also in their concave shape (cross-sectional area), mostly due to the presence of a dorsal groove. In addition, we find an increase in cortical medulla and the presence of transversal septa that act as internal support, as previously noted [2]. This earlier work suggested that the existence of a groove in these spiny hairs increased surface area, which, in addition to specific ultrastructural cuticle scales, likely played an important thermoregulatory (heat loss) and water condensation function in tropical and subtropical species. Moreover, the effect of increasing cross-sectional area and especially the cortical layer in reducing flexibility has been long recognized, at least in plant species [37]. Development of such grooves reduces torsion along the spine, an important characteristic associated with the effectiveness of spine penetration. These characteristics may explain the greater robustness and concavity as well as the relative increase in cortical area of spiny hairs compared to guard hairs. Moreover, these traits are required for the function of this type of spine. For example, in the Old World porcupine (*A. macrourus*), in which the spines have clear protective function [38], the tip is inserted into the body of predators during defense based on the axial compression load. In some cases, the entire retraction of the spine occurs. Alternatively, the tip, which is shorter and presents a smaller cross-section (when compared to the middle or base of the spine) breaks and remains inside the body of predators [2].

A second evolutionary lineage corresponds to the circular section of spines present only in the New World porcupines (Erethizontidae). These circular and stiff spines have a morphology that maximizes the elastic buckling compared to the other five spiny rodent lineages studied. This biomechanical property is due largely to the small cross section relative to length, the lack of cortex, and the abundance of soft alveolar medulla that is continuous and without septa [39]. The penetration of this type of spine in the predator/opponent, which is deeper (compared to the previous case) is indeed initially associated with axial compression by buckling. Accordingly, the spine breaks but generally in the proximal region instead of the tip and remains in the body of a predator [2]. The low-tension values indicate flexibility of this particular circular hair shape, in which stable and constant values of tension represent the greatest deformation without strain and breakage, making it ideal for predator defense – the spine is inserted without it breaking.

Thus, a “spiny hair phenotype” can be achieved in multiple ways. First, the concave hair of Old World porcupines likely represents a distinct evolutionary path from the New World porcupines, even if facing similar ecological challenges. While American porcupines have hairs with greater flexibility (in which hair tension values do not exceed 40 MPa until 30-40% of deformation), hair from the Old World porcupines have increased tension (around 100 MPa in the first 30% of deformation). Second, and contrary to our expectations, the two independent morphologies associated with spiny hairs of wild rodents are distinct from that of East Asian human populations [27, 28].

As the detailed morphological shape and function of spiny hair observed in wild rodents – the grooved and circular forms – were distinct from that of Asian humans, it was perhaps not surprising that the EDAR 370A mutation was not present in any species surveyed, even in the New World porcupines that are morphologically most similar to Asian hair. We did, however, find substitutions in the first and third position of this 370 codon that resulted in one amino acid change (Val^370^Iso), but this variant was not associated with hair morphology. Specifically, this 370Iso mutation was present in both spiny and non-spiny sister species (kangaroo rat and spiny pocket mouse), clearly consistent with their close evolutionary relationship and not associated with hair type. When we look across mammalian orders, we found that 370Iso is also present in the lagomorph pika, the primate mouse lemur, and two lineages of marsupials, tasmanian devil and wallaby opossum, indicating that this codon has not been conserved over evolutionary time. A second amino acid change (L^422^V) was observed only in echimyid rodents. We were able to collect sequence data at this 422 site from only a few taxa (all non-spiny controls) because of its proximity to the end of the EDAR death domain, which limits our ability to compare genotypes in other spiny rodents. However, when we again look across deeper divergence, we found that guinea pigs (a Hystricomorpha) also have the 422Val variant. Moreover, we found this mutation in two other Hystricomorpha taxa (*Euryzygomatomys spinosus* and *Agouti paca*) (unpublished data). While all these hystricomorph species are characterized by thick or wiry hairs, testing if this substitution affects hair concavity will require further morphological and genetic experiments.

Although strongly associated with EDAR, variation in hair thickness among human populations cannot be explained exclusively by the V370A polymorphism [28], suggesting that other genetic variants are associated with hair morphology. Genetic variants that affect the shape of hairs or cross-sectional area have yet to be identified and functionally validated. However, in recent work, Fujimoto et al. [40] examined an additional ten candidate genes: *Lef1*, *Msx2*, *Dll1*, *Egfr*, *Cutl1*, *Notch1*, *Fgfr2*, *Krt6irs*, *Gpc5*, *Akt1*, *Myo5a*, *Tgm3*, *Eda2R* and *Eda*. A significant association was observed between a G/T SNP (rs4752566) in the intron nine of the gene *Fgfr2* and human hair morphology, which showed the strongest association with cross-sectional area and diameter size. While still preliminary, these results suggest that other genes are likely involved in hair differentiation. Thus, either different mutations in the *Edar* gene or mutations in other genes likely contribute to the evolution of spiny-hair phenotypes in natural populations of mammals, including the rodents surveyed in this study.

In conclusion, our results show that, at a morphological level, there are at least two evolutionary paths that result in a spiny-hair phenotype in wild rodents. Compared to an elliptical shape of hairs in non-spiny rodents, we found hairs that were grooved in cross-section (in the bristle mouse, spiny pocket mouse, Southern African spiny mouse, tree rats and Asiatic brush-tailed porcupine) and an almost perfectly cylindrical form, present in the New World porcupines. These morphological types, in turn, lead to measurable differences in tensiometric properties, between each other and relative to non-spiny control mice as well as human hair. In addition, while a polymorphism in the *Edar* protein (V^370^A) was associated with straighter and thicker hairs in human populations, in the lineages of rodents we examined, we did not find an association between this variant of the *EDAR* protein and hair phenotype. Thus, the non-synonymous substitution in the Edar previously associated with “spiny” hair morphology in humans does not play a role in modifying rodent hairs in this study, suggesting different mutations in *Edar* and/or other genes are responsible for variation in the spiny hair phenotypes observed in wild rodent species.

## Acknowledgements

We thank to Judith Chupasko (MCZ, Harvard University), Chris Conroy (MVZ, University of California, Berkeley), Ingrid Rochon (Smithsonian Institution, National Museum of Natural History) and Bruce Patterson (Field Museum of Natural History) for kindly providing DNA samples and rodent hairs. We also thank the Electron Microscopy Center at UFRGS for use of the micrograph equipment.

## Supporting Information

**S1 Fig.**
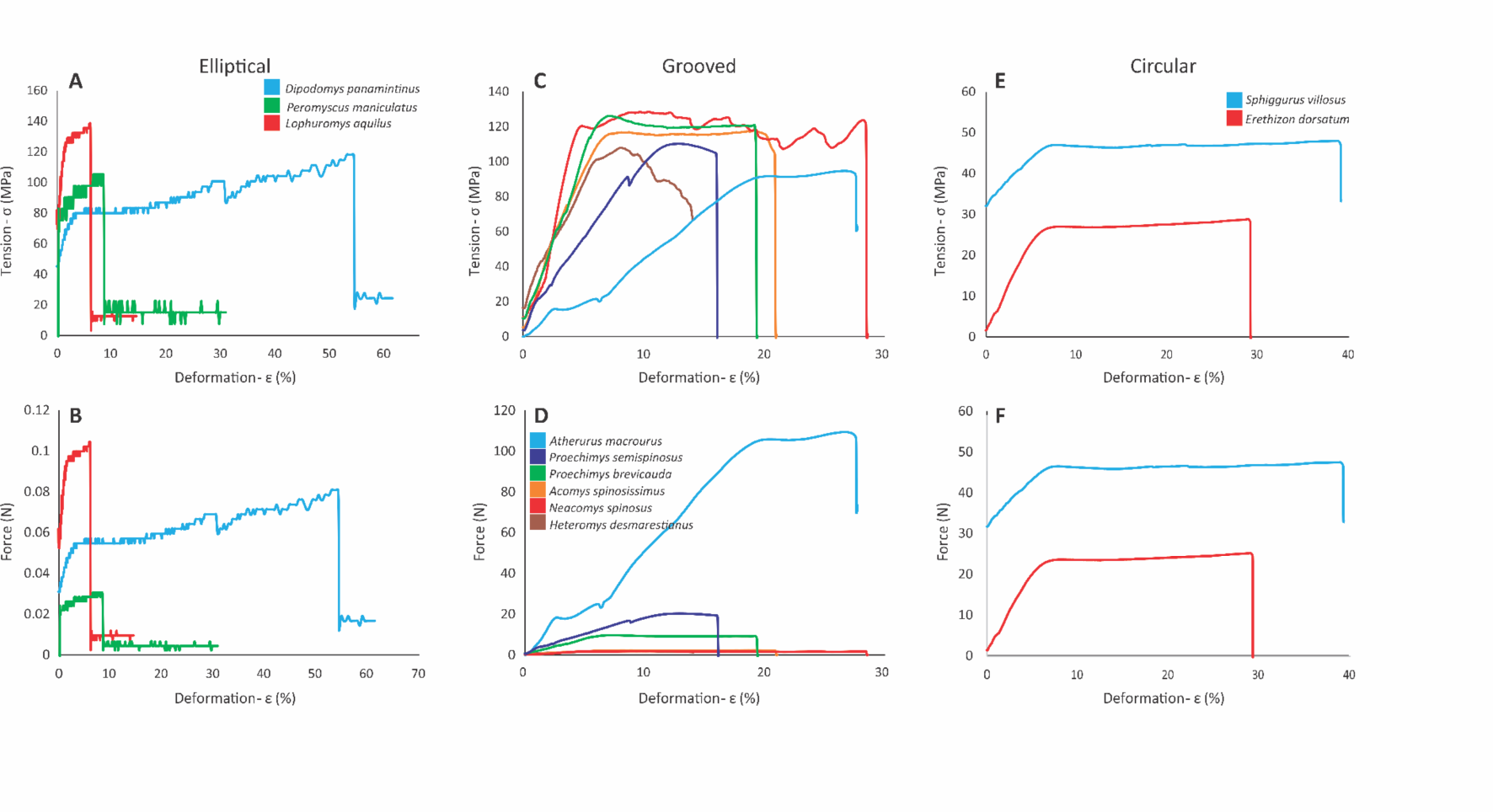
Deformation curves in relation to tension (A, C, E) and force (B, D, F) for hairs of eleven rodent species, split according to the cross-section shape of the hair.

**S2 Fig.**
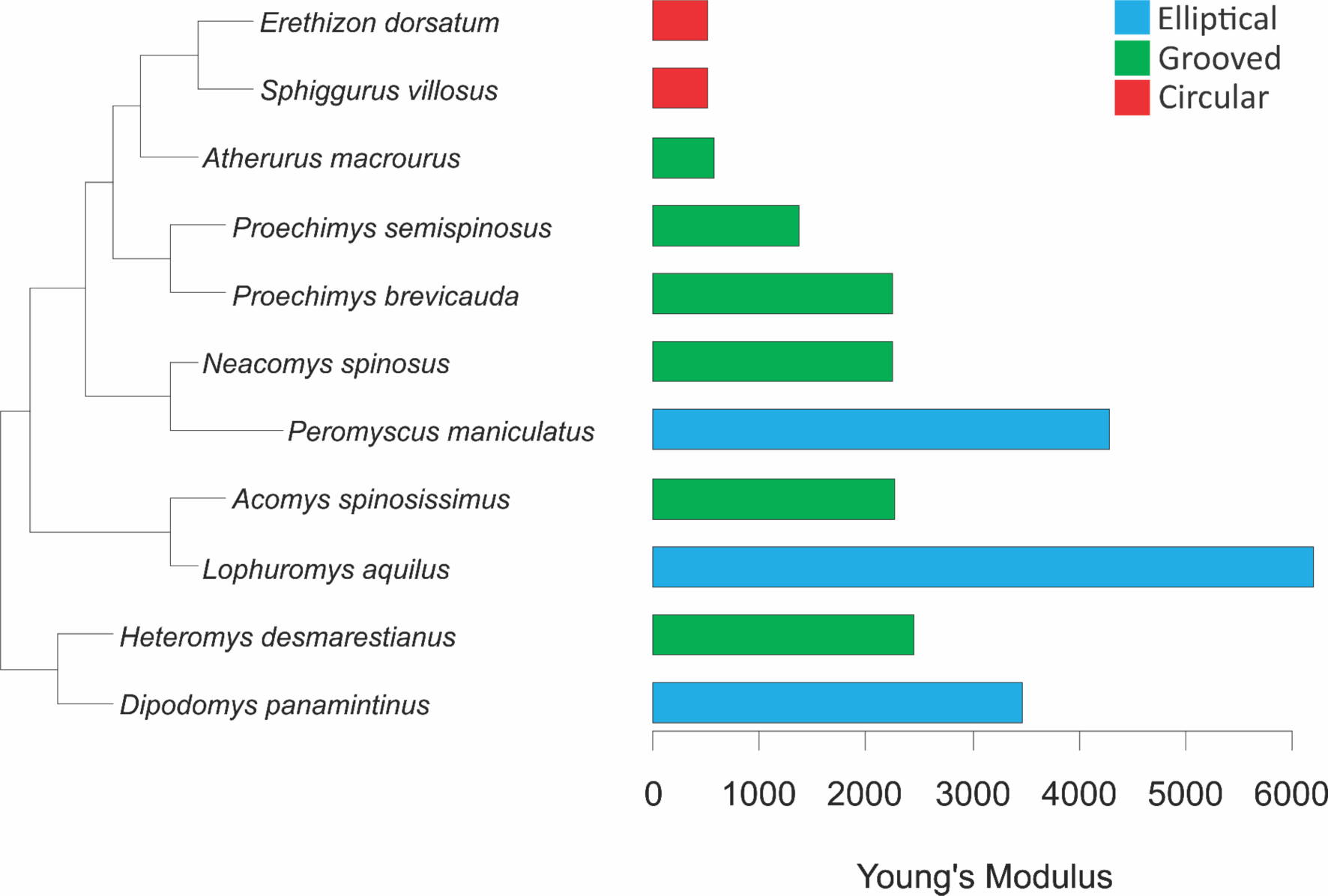
Young’s modulus for the linear portion of hair deformation curves (Fig 5A) for eleven rodent species. The modulus was calculated as E= σ/ε for the initial portion of each deformation curve (at 3% of deformation).

